# Exploring the Single-Cell Dynamics of FOXM1 Under Cell Cycle Perturbations

**DOI:** 10.1101/2024.07.27.605093

**Authors:** Tooba Jawwad, Maliwan Kamkaew, Kriengkrai Phongkitkarun, Porncheera Chusorn, Somponnat Sampattavanich

## Abstract

The cell cycle is crucial for maintaining normal cellular functions and preventing replication errors. FOXM1, a key transcription factor, plays a pivotal role in regulating cell cycle progression and is implicated in various physiological and pathological processes, including cancers like liver, prostate, breast, lung, and colon cancer. Despite previous research, our understanding of FOXM1 dynamics under different cell cycle perturbations and its connection to heterogeneous cell fate decisions remains limited. In this study, we investigated FOXM1 behavior in individual cells exposed to various perturbagens. We found that different drugs induce diverse responses due to heterogeneous FOXM1 dynamics at the single-cell level. Single-cell analysis identified six distinct cellular phenotypes: on-time cytokinesis, cytokinesis delay, cell cycle delay, G1 arrest, G2 arrest, and cell death, observed across different drug types and doses. Specifically, treatments with PLK1, CDK1, CDK1/2, and Aurora kinase inhibitors revealed varied FOXM1 dynamics leading to heterogeneous cellular outcomes. Our findings affirm that FOXM1 dynamics are pivotal in determining cellular outcomes, independent of the specific inhibitor employed. Our results gave insights into how FOXM1 dynamics contribute to cell cycle fate decisions, especially under different cell-cycle perturbations.

## INTRODUCTION

Understanding the cell cycle is crucial for comprehending how cells grow, divide, and ensure the continuity of life. Its precise regulation is vital for maintaining both the development and the repair of tissues in multicellular organisms. Cell cycle dysregulation has been implicated in various diseases, most notably cancer, where uncontrolled cell proliferation and genomic instability are among major cancer hallmarks (1). In the intricate network of cell cycle regulation, the transcription factor FOXM1 emerges as a key player (2).

FOXM1, a member of the FOX family of transcription factors, plays a crucial role in cell cycle regulation (3). It promotes cell cycle progression by driving the expression of genes necessary for G1/S transition, DNA repair, and angiogenesis (4). FOXM1 interacts with key regulatory proteins like p53, Rb, and CDKs (CDK2-cyclin E, CDK1-cyclin B) involved in cell cycle checkpoints (5, 6). FOXM1 is regulated by the cell cycle, but itself can also affect cell cycle progression. For example, FOXM1 is subject to tight regulation by signaling pathways Aurora-A and PLK1 which regulate FOXM1 promoter activity, forming a positive feedback loop that enhances FOXM1 and their own expression. (7, 8). The tight regulation of FOXM1 is essential for ensuring the accurate timing and orderly progression of key cell cycle events (9).

FOXM1’s crucial role in cell cycle regulation is linked to various pathological conditions, especially oncogenesis. Aberrant FOXM1 expression correlates with increased cell proliferation, apoptosis resistance, and tumor progression, as documented in multiple studies (2, 3, 9-12). Specifically, FOXM1 malfunctions are often tied to G1 checkpoint anomalies, leading to inappropriate expression timing and levels driven by the loss of regulatory controls (6, 13). This results in disrupted cell cycle dynamics and, potentially, genomic instability (4, 10, 13). Furthermore, studies have demonstrated that such dysregulation can induce cellular transformation and a neoplastic phenotype, underscoring FOXM1’s role in cell cycle regulation breaches and its impact on tumorigenesis (14-16).

Our understanding of FOXM1’s role in cell cycle progression and disease pathogenesis, traditionally informed by bulk measurements, often overlooks cellular heterogeneity in FOXM1 expression. Recent advancements in single-cell signaling dynamics via live microscopy, such as those highlighted by Spencer et al. (2013) studying mitotic exit populations controlled by p21 with a CDK2 reporter, offer insights into this heterogeneity (17). These tools are crucial for comprehending FOXM1 dynamics, particularly where the cell cycle is perturbed. Insights gained from studying FOXM1’s role under cell cycle perturbations can identify the relationship between the FOXM1 dynamics and cellular phenotypes.

Our research aims to address this knowledge gap by deploying live microscopic imaging of cells with a FOXM1 reporter. This approach allows us to examine the dynamical behavior of FOXM1 at the single-cell resolution and to investigate the corresponding heterogeneous cell fate decisions under different cell cycle perturbations. We aim to comprehend the diverse patterns of FOXM1 dynamics and investigate how they are correlated with the diversity of cell fate determinations. Consequently, our work plays a pivotal role in offering a thorough perspective on FOXM1 dynamics, potentially revealing the temporal intricacies of FOXM1 under different cell cycle conditions.

## MATERIALS AND METHODS

### Cell Culture

Human mammary epithelial MCF10A stably expressing FOXM1-mVenus-H2B-mCherry cells were cultured as previously described (18). MCF10A full growth media consisted of Dulbecco’s modified Eagle’s medium (DMEM)/F12 (Life technologies 11330), supplemented with 5% horse serum (Life Technologies), EGF (20ng/ml, Preprotech), bovine insulin (10 ug/ul, Sigma), hydrocortisone (0.5 ug/ml), cholera toxin (100 ng/ml, Sigma) and penicillin (50 U/ml) and streptomycin (50 ug/ml). Serum starvation for the MCF10A cell line was done with MCF10A full growth media without horse serum, EGF, and bovine insulin supplementation.

### Cell Line Construction

The FOXM1 sensor was constructed by Gibson assembly, combining fragments of FOXM1B (NM_202003) fused with mVenus and a Histone 2B fragment fused with mCherry into a PiggyBac vector. This polycistronic construct, pPB-FoxM1-mVenus-P2A-H2B-mCherry, along with a hyPBase transposase vector, was co-transfected into MCF10A cells using Lipofectamine LTX. Transfected cells were cultured for at least two weeks and sorted using BD FACSAria II to obtain a pure population expressing the fluorescent reporter. Cells were maintained in MCF10A full-growth media consisting of DMEM/F-12 supplemented with 5% horse serum, EGF (20 ng/ml), insulin (10 μg/ml), hydrocortisone (0.5 μg/ml), cholera toxin (100 ng/ml), penicillin (50 U/ml), and streptomycin (50 μg/ml) as previously described (18).

### Western Blotting

Total proteins were extracted with RIPA buffer containing protease inhibitors, separated on 7% SDS PAGE gels and transferred into the PVDF membranes (GE). Then, incubated at 4°C for 24 h with the primary antibody (Rabit-anti FOXM1 #D12D5, Cell Signaling) at a ratio of 1:1000. Primary antibodies were visualized using a secondary antibody conjugated to IRDye fluorophores (700 nm) (Li-Cor) in Odyssey PBS blocking buffer at a ratio of 1:2500 for 1 h. Membranes were scanned using an Odyssey scanner.

### Luciferase Assay

MCF10A carrying FOXM1-mVenus-H2B-mCherry reporter was plated in a 96-well plate for 24 h in advance at 90% confluency. Cells were transfected with the human FOXM1 promoter (19) and Renilla (pRL-TK, Promega) as an internal transfection control using Lipofectamine LTX (#15338100, Invitrogen) according to the manufacturer’s instructions for 24 h. Cells were harvested and then measured firefly/Renilla luciferase activity using Dual-Glo® Luciferase assay (#E2920, Promega).

### Inhibitors

A443654 (AKT; SelleckChem), Palbociclib (CDK4/6; SelleckChem), Abemaciclib (CDK4/6; SelleckChem), RO3306 (CDK1; Santa Cruz), K03861 (CDK2; (SelleckChem), BI2536 (PLK1; MedChem Express), Volasertib (PLK1; MedChem Express), Tozasertib (Aurora Kinase; MedChem Express), Danusertib (Aurora Kinase; MedChem Express), Rabusertib (CHK1; MedChem Express), Prexasertib (CHK1; MedChem Express), Berzosertib (ATR; MedChem Express), Bay1895344 Hydrochloride (ATR; MedChem Express), Ceralasertib (ATR; SelleckChem), BMS265264 (CDK1/2; MedChem Express), and CDK1/2 III (CDK1/2; Santa Cruz) were dissolved in DMSO (Sigma) with stock concentration at 10 mM. All stock was kept at -80 degrees Celsius for long-term storage. The final concentration used in this study was 0 µM, 0.3µM, 0.6µM, 1.25µM, 2.5µM, 5µM and 10µM.

### Time-lapse Microscopy

MCF10A cells expressing FOXM1-mVenus-H2B-mCherry reporter were seeded at 70% confluency in 96-well plates for 24 h. After three washes with DMEM/F12 base media, cells were starved in MCF10A-starve media for 24 h synchronize in G1 phase. Following synchronization, cells were treated with various concentrations (0 to 10 µM) of cell cycle inhibitors before serum replenishment. Images were captured every 10 mins over 48 h using YFP (Ex: 508/24, Em: 540/21) and RFP (Ex: 575/22, Em: 632/60) filters after replacing starved media with full-growth media containing inhibitors. A 0.1% v/v diluted DMSO served as a negative control for compound treatments.

### Immunofluorescence Staining

Cells were fixed with 2% paraformaldehyde for 15 minutes and stained with 1:1000 Hoechst 33342 for 30 minutes. Imaging for Hoechst, mCherry, and mVenus was conducted using the CLS high-content system. After imaging, cells were permeabilized with pure methanol for 15 minutes. Fluorescence was bleached according to the CyCIF protocol (20), to enable subsequent staining with EdU and DAPI, followed by re-imaging using CLS high-content imaging. Images from both imaging sessions were aligned and quantified using CellProfiler.

### Image Processing and Analysis

Single-cell fluorescence quantification was performed using CellProfiler. Blank well images provided illumination and background immunofluorescence intensities for flatfield correction and background subtraction. Nuclei were segmented based on DAPI signal, with cytoplasm identified as an extended region 5 pixels beyond the nuclei boundary. Mean and integrated fluorescence intensities for mCherry, mVenus, DAPI, and EdU were measured in both nucleus and cytoplasm compartments.

### Cell Cycle Segmentation

All cell cycle segmentation was done automatically by using 2 criteria. First, the EdU intensity of all treatment conditions was collected, and then a cut-off between negative and positive EdU staining was generated by using a bimodal distribution. Then, similar procedure was applied to DAPI. These two cut-offs segment the cell cycle into 4 segments: the G1 phase (low DAPI and low EdU intensity), S phase (low DAPI and low EdU intensity); the early S phase (low DAPI and high EdU); the late S phase (high DAPI; and high EdU); and then finally, G2/M phase (high DAPI with low EdU).

### Cell Tracking

Initially, the raw images underwent processing using the cell profiler to perform background subtraction and illumination correction for raw images. FUCCITrack software was then used for the single-cell tracking as previously described (21).

### Statistical Analysis

All statistical analyses were performed utilizing GraphPad Prism® software. Data are shown as mean ± standard deviation (SD). Statistical analysis was done with respect to the control using an unpaired two-tailed Student’s t-test.

## RESULTS

### Construction and characterization of live-cell reporter for FOXM1 translocation dynamics

Our objective was to develop a FOXM1 reporter system to observe its translocation from the cytosol to the nucleus under various cell cycle perturbation conditions. We used the full-length FOXM1B cDNA to produce this reporter as our template. The FOXM1-mVenus reporter was constitutively expressed under the control of the chicken beta-actin promoter. To enable the monitoring of the nuclear compartment, we co-expressed the FOXM1-mVenus reporter and the H2B-mCherry nuclear reporter into a polycistronic construct, separated by the ribosomal skipping P2A sequence (Fig. 1a). As expected, we observed a correlated expression of the FOXM1-mVenus and the H2B-mCherry reporter (Fig. S1a). This concurrent expression of both enabled us to monitor continuously and positively sort cells with stable expression of the FOXM1-mVenus based on the H2B intensity.

**Figure 1.**
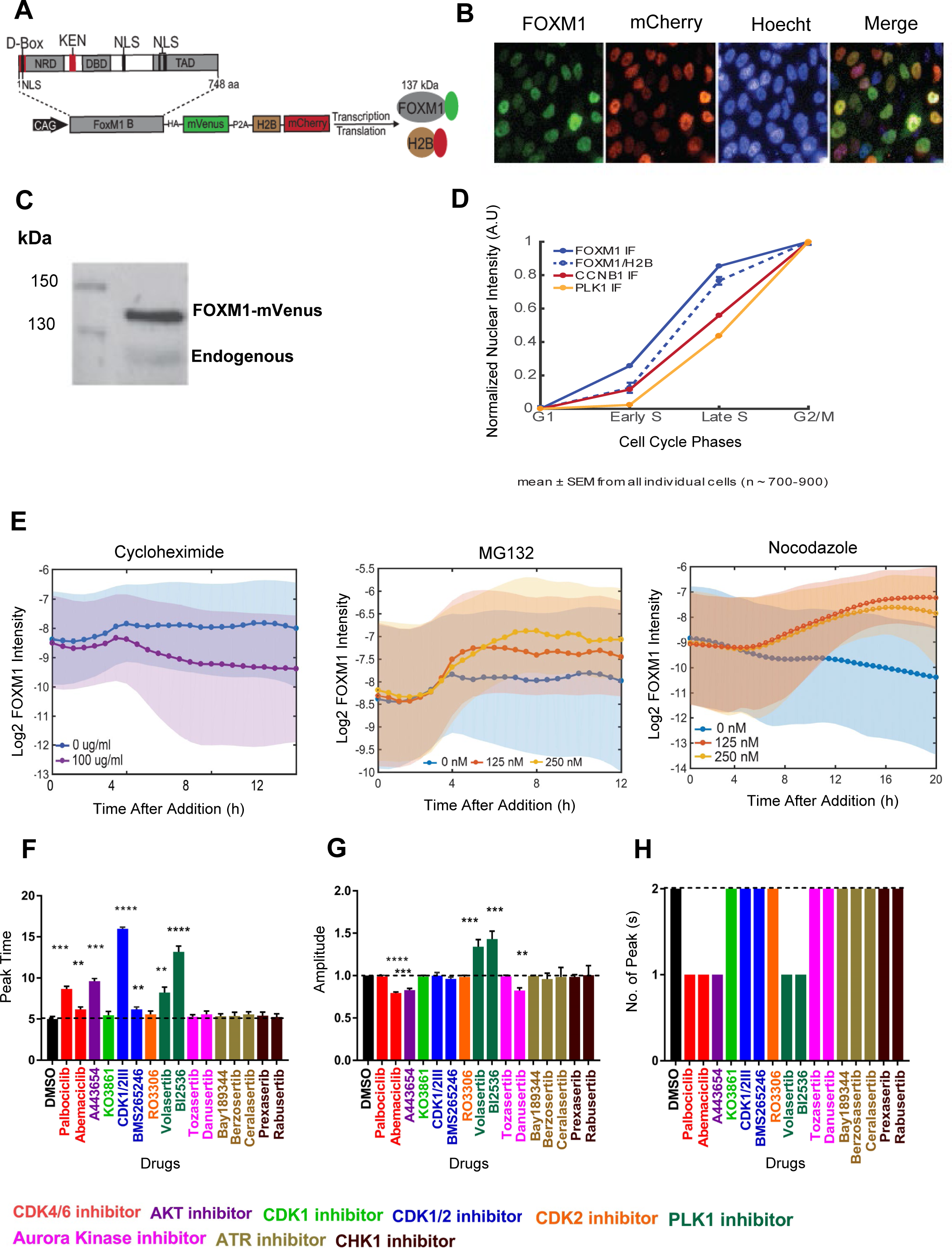
FOXM1 reporter faithfully reports FOXM1 dynamics. (**A**) FOXM1 reporter engineered by fusing FOXM1 B with mVenus, linked with Histone 2B-mCherry by P2A peptide. (**B**) Fluorescent microscopic images of MCF10A-FOXM1 reporter at different channels (FOXM1-mVenus, H2B-mCherry, Hoecht, and Merge) from live imaging. (**C**) Western blot of the FOXM1 reporter and endogenous FOXM1. (**D**) Comparison of endogenous FOXM1 (blue line), FOXM1 reporter (dotted blue line), CCNB1 (red line), and PLK1 (yellow line) at different cell cycle phases. (**E**) Changes of FOXM1 reporter when exposed to Cycloheximide (0 and 100μg/ml), Proteasome inhibitor MG132 (0, 125, and 250 nM), and Nocodazole (0, 125, and 250 nM). (**F-H**) Comparison of FOXM1 dynamics after treatment with 16 cell cycle perturbagens at 5μM for Tozasertib, RO3306, and CDK1/2III, 2.5μM Danusertib, A443654, and KO3861 and 10μM for the rest (n=3 per each condition). Bulk FOXM1 dynamics were compared based on the onset time of the first peak (F), amplitude of the first peak (G), and number of peaks (H).

We first examined the nuclear localization of the FOXM1-mVenus reporter, which predominantly localized in the nucleus compared to H2B-mCherry and Hoechst signals (Fig. 1b). Transcriptional activity of FOXM1-mVenus was 1.8-fold higher than endogenous FOXM1 in parental MCF10A cells (Fig. S1b), with its abundance approximately 2-fold higher based on western blot analysis (Fig. 1c). Increased FOXM1 abundance led to a slight 0.5-fold increase in genes such as CDC25B, NEK2, and KNSTRN, without significant changes in other FOXM1 targets (Fig. S1c). Importantly, cyclin B1 abundance showed no significant differences between reporter-expressing cells and the parental MCF10A cell line in western blot analysis (Fig. S1d), suggesting the FOXM1 reporter accurately monitors FOXM1 translocation dynamics.

FOXM1 is also well known as one of the important cell-cycle proteins. The level of FOXM1 increases during the S phase and maximizes in the G2/M phase. To validate its cell-cycle-dependent behavior, we utilized Cyclic Immunofluorescence (CycIF) with our FOXM1-mVenus reporter. This technique allowed us to monitor FOXM1-mVenus dynamics alongside DAPI and Edu staining for cell-cycle phase annotation (Fig. S1e) (20). Analysis revealed that FOXM1-mVenus changes throughout the cell cycle mirrored those of endogenous FOXM1 (Fig. 1d). We also tracked Cyclin B1 and PLK1, both known FOXM1 interactors, observing their increased levels during S and G2/M phases (Fig. 1d). These findings align with Cyclin B1’s G2 accumulation and PLK1’s dynamics through the cell cycle stages (22, 23). Importantly, our reporter did not alter FOXM1 target protein dynamics.

We next examined changes in the FOXM1 reporter under known perturbagens.; Cycloheximide, MG132, and Nocodazole (Fig. 1e, Fig. S1f). Cycloheximide reduced FOXM1 reporter abundance (Fig. 1e-left panel), with a half-life of ∼0.5 hours (Fig. S1g), confirming APC/C E3 ubiquitin ligase-mediated degradation, akin to endogenous FOXM1 (24). Conversely, MG132 stabilized FOXM1-mVenus for up to 12 hours (Fig. 1e-middle panel). FOXM1 reporter intensity gradually increased under Nocodazole, reaching stability by 20 hours, aligning with M phase arrest (25). We concluded that our carefully sorted pool of the MCF10A cell line with the stable expression of the FOXM1-mVenus reporter effectively delineates the temporal dynamics of the endogenous FOXM1.

### Quantitative analysis of FOXM1 dynamics in response to cell cycle perturbagens

Taking advantage of our reporter, we investigated FOXM1 dynamics across different cell cycle phases under various perturbagens over 48 hours. Cells were initially serum-starved for 24 hours, treated with inhibitors or DMSO for 30 minutes, and then imaged every 10 minutes upon release into full growth media to monitor FOXM1 dynamics (Fig. S2a). In the DMSO control, FOXM1 nuclear intensity initially peaked at 4 hours post-release, followed by a subsequent decrease (Fig. S2b). We further analyzed FOXM1-mVenus dynamics in response to all inhibitors, focusing on parameters such as peak time, number of peaks, and first peak amplitude (Fig. S2c).

Notably, a statistically significant increase in the peak time of FOXM1 was observed under treatment with CDK4/6 (Palbociclib and Abemaciclib), CDK1/2 (CDK1/2III and BMS 265246), ATR (Bay189344 Hydrochloride, Berzosertib, and Ceralasertib), and PLK1 (BI2536 and Volasertib) inhibitors (Fig. 1f). Additionally, the amplitude of FOXM1 was significantly reduced with CDK4/6 (Abemaciclib), AKT (A443654), and Aurora Kinase (Danusertib) inhibitors, while a contrasting increase was observed in response to the PLK1 (BI 2536 and Volasertib) inhibitors (Fig. 1g). Interestingly, only a single peak of FOXM1 was observed with CDK4/6, AKT, and PLK1 inhibitors (Fig. 1h), aligning with prior studies indicating G1 arrest induced by Palbociclib and Abemaciclib (26, 27), and G1/S transition inhibition by AKT (28). These findings unravel the nuanced and inhibitor-specific modulation of FOXM1 dynamics.

### Characterizing the dynamics of FOXM1 reporter using functional principal component analysis

To distinguish the differential responses of FOXM1 dynamics systematically, we utilized functional principal component analysis (fPCA) to decompose FOXM1 nuclear signals from cells exposed to 16 inhibitors over 48 hours (t = 0 to 48 h). The analysis captured diverse FOXM1 intensity patterns effectively using the primary harmonic functions, fPC1 and fPC2, which explained over 95% of the variance (Fig. 2a, b-upper panel). fPC1 depicted sustained FOXM1 activity increasing gradually from t = 1 to 48 h, while fPC2 showed a transient pattern with an initial rise and subsequent decline from t = 1 to 48 h (Fig. 2b-lower panel).

**Figure 2.**
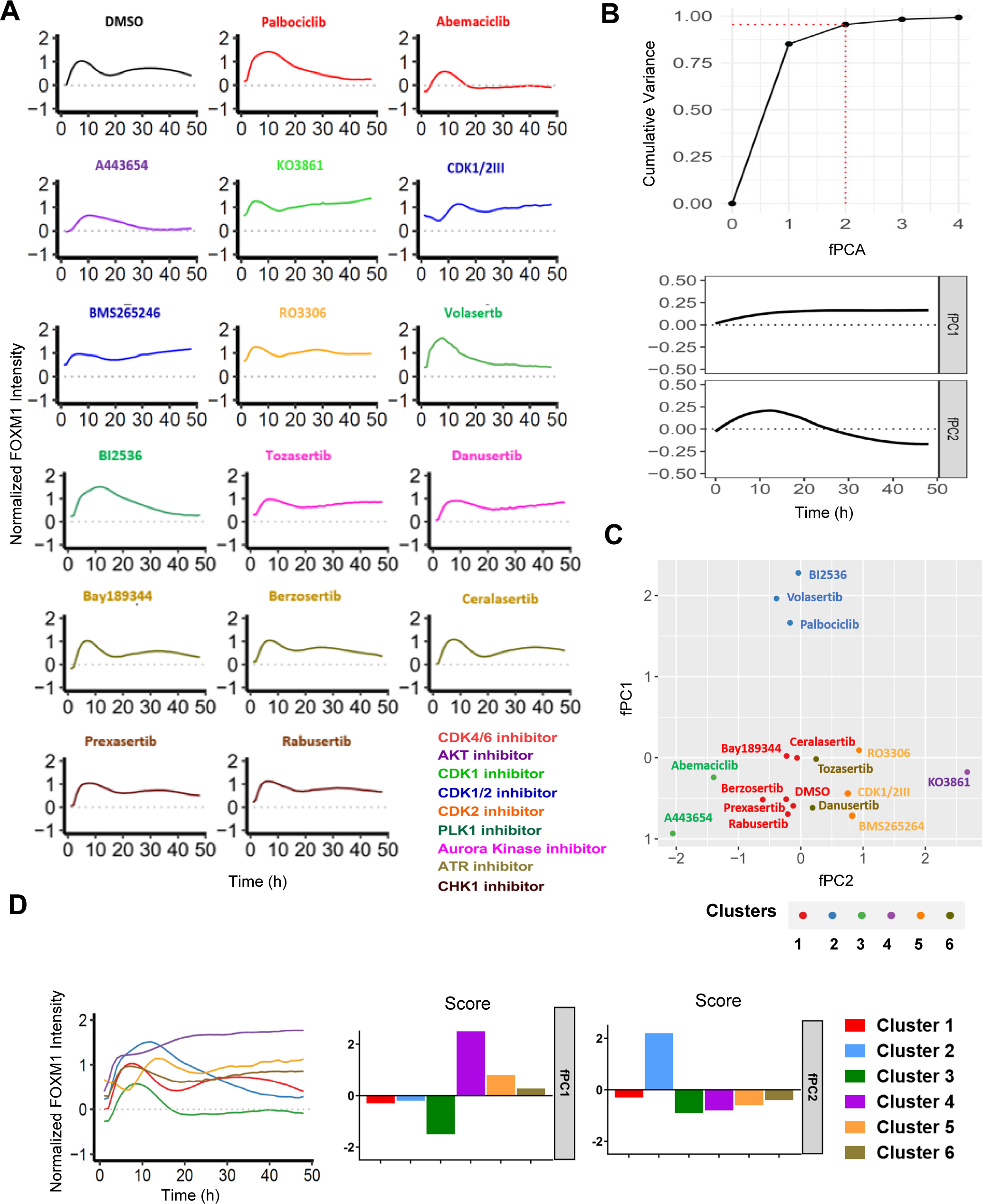
Functional principal component analysis (fPCA) of FOXM1 dynamics under different cell cycle perturbagens. (**A**) Normalized FOXM1 activity after treatment with 16 cell cycle perturbagens at 5μM for Tozasertib, RO3306, and CDK1/2III, 2.5μM Danusertib, A443654, and KO3861 and 10μM for the rest, plotted over a 48 h. (**B**) Cumulative variance explained by FOXM1 dynamics across fPCA scores (t=0 to 48 h). Upper: The first two fPCA scores collectively explain over 95% of the total variance (red dotted line) across all treatments. Lower: fPC1 captures a gradual increase in FOXM1 activity from 0-10 h followed by sustained levels (11-48 h), while fPC2 shows an initial surge in FOXM1 activity (0-10 h) followed by a decrease (11-48 h). fPC1 and fPC2 together account for 85% and 90% of the total variance, respectively. (**C**) The fPC1-vs-fPC2 score plot of all 16 cell cycle perturbagens reveals clustering into 6 subgroups by KNN algorithm: cluster 1 (red), cluster 2 (blue), cluster 3 (green), cluster 4 (purple), cluster 5 (orange), and cluster 6 (olive). (**D**) Comparison of averaged FOXM1 dynamics across 6 subgroups. Upper: Temporal trajectories of FOXM1 dynamics averaged for all 6 cluster. Lower: Comparison of fPC1 and fPC2 scores among the 6 clusters.

In the fPC1-fPC2 space, inhibitors clustered into six distinct groups based on KNN clustering algorithm and their fPC scores (Fig. 2c). For instance, DMSO, ATR and CHK1 inhibitors formed Cluster 1 due to their similar dynamic patterns and fPC scores (Fig. 2c, d). PLK1 inhibitors clustered with the CDK4/6 inhibitor Palbociclib in Cluster 2, despite having different molecular targets (Fig. 2c). While Cluster 4 comprises of CDK2 inhibitor, and Cluster 5 consists of a CDK1 inhibitor and CDK1/2 inhibitors (Fig. 2c). Aurora kinase inhibitors grouped together in Cluster 6 (Fig. 2c). Notably, CDK4/6 inhibitor Abemaciclib showed unique dynamics with a high negative fPC1 score and a low fPC2 score, placing it in Cluster 3 with AKT inhibitor, distinct from Palbociclib, which had a high positive fPC2 score (Fig. 2d-lower panel).

This analysis highlights distinct clusters of FOXM1 dynamics induced by the 16 inhibitors, revealing unexpected groupings that underscore the specificity and complexity of inhibitor effects on FOXM1 dynamics, such as Palbociclib, Volasertib, and BI2536. This unexpected grouping underscores the specificity and complexity of different inhibitors in FOXM1 dynamics. These observations are consistent with our previous findings in Fig. 1f and 1h and Supplementary Fig. 2b, where the dynamic pattern of FOXM1 influenced by Palbociclib, Volasertib, and BI2536 is similar.

### Cell fate decisions correspond with FOXM1 dynamics across different cell cycle perturbagens

We hypothesized that drugs inducing similar FOXM1 activity dynamics would lead to consistent phenotypic outcomes. After 24 h of treatment, cells were fixed, photo-bleached, and stained with DAPI and Edu for cell cycle stage identification, utilizing DAPI for DNA content and image alignment (Fig. 3a). ATR and CHK1 inhibitors showed FOXM1 dynamics similar to DMSO (cluster 1), with corresponding cell cycle distributions (Fig. 3b). PLK1 inhibitors and Palbociclib shared FOXM1 dynamics as cluster 2 perturbagens and exhibited a dominant G1-arrested phenotype (Fig. 3b). Similarly, Abemaciclib and A443654 displayed similar FOXM1 dynamics (cluster 3 in Fig. 2c, d) and phenotypic outcomes, including a G1-arrested phenotype with a subset in G2/M (Fig. 3b). CDK1/2 and CDK1 inhibitors (cluster 5) and Aurora kinase inhibitors (cluster 6) induced G2/M arrest, with slight variations in phase distribution (Fig. 3b). These findings underscore the correlation between FOXM1 dynamics and cell cycle phenotypic outcomes.

**Figure 3.**
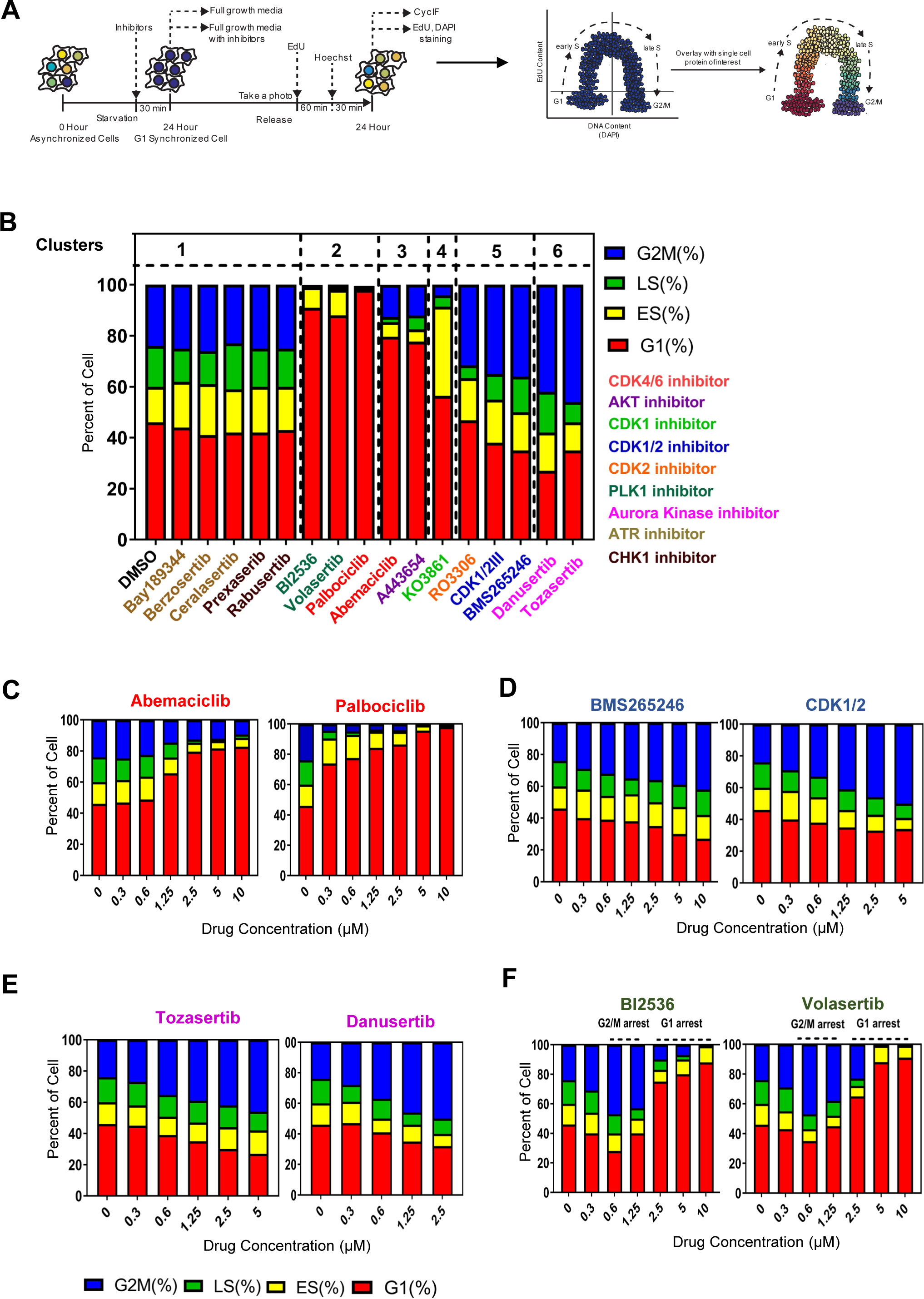
Comparison of phenotypic outcomes between the 6 different FOXM1 dynamic clusters. (**A**) Workflow for annotating cell cycle stages post-live imaging of the FOXM1 reporter: Following serum-starve synchronization and treatment with 16 cell cycle inhibitors for 24 h, cells were fixed and restained with EdU and DAPI using the CycIF protocol. EdU incorporation and DNA content were utilized to annotate cell cycle stages: G1, early S, late S, and G2/M. (**B**) Cell cycle distribution of MCF10A cells expressing the FOXM1 reporter, grouped according to drug clusters based on fPCA scores (n=3 per condition). Drug doses: 5μM for Tozasertib, RO3306, and CDK1/2III; 2.5μM for Danusertib, A443654, and KO3861; 10μM for the remaining drugs. (**C-F**) Cell cycle distribution changes with altering drug concentration. n=3 per condition.

We next asked whether drugs of the same class exhibit dose-dependent phenotypic differences. For this purpose, we analyzed outcomes from various drug targets. Interestingly, Palbociclib and Abemaciclib, both CDK4/6 inhibitors, showed distinct responses. Abemaciclib induced G1 arrest at higher doses (10µM to 1.25µM) and reverted to a control-like phenotype at lower doses, whereas Palbociclib consistently induced pronounced G1 arrest across all concentrations (Fig. 3c). Similarly, CDK1/2 inhibitors exhibited a dose-response relationship, with decreasing proportions of G2/M arrest observed at lower concentrations (0.6µM) (Fig. 3d). Aurora kinase inhibitors induced G2/M arrest-like phenotypes at varying doses (Fig. 3e). PLK1 inhibitors induced G1 arrest at higher doses (10 to 2.5µM) and G2/M arrest at lower doses (1.25 and 0.6µM) (Fig. 3f).

Observing varied phenotypes at different concentrations prompted us to explore the role of dosage in influencing cellular outcomes. We exposed MCF10-mVenus reporter cells to 16 inhibitors across concentrations from 0 µM to 10 µM. Figure 4a illustrates how FOXM1 dynamics are modulated by dosage, suggesting that FOXM1 signals, rather than just the inhibitors themselves, drive specific cellular responses (G1 arrest, G2/M arrest, or similar to DMSO control). A bimodal FOXM1 peak typically correlates with the DMSO control cell cycle distribution (Fig. 4b, c). Delayed FOXM1 peaks (Fig. 4b-upper panel) are associated with G1 arrest phenotypes (Fig. 4b-lower panel), while sustained FOXM1 increases at later stages (Fig. 4c-upper panel) correspond to G2/M arrest (Fig. 4c-lower panel). We concluded that FOXM1 dynamics (signals) are the critical determinant of cell fate decisions (response).

**Figure 4.**
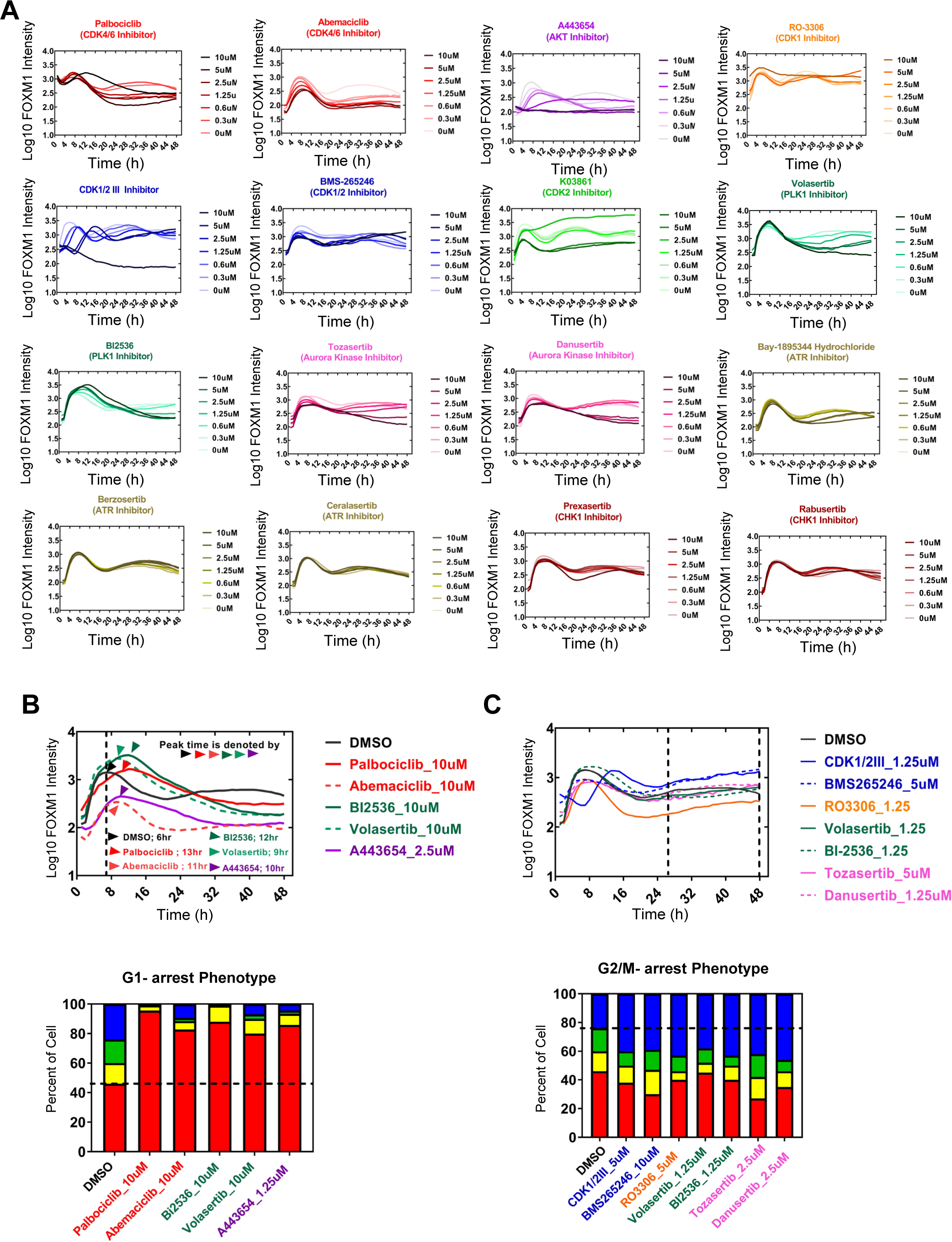
Prediction of cell fate decision by FOXM1 dynamics from different drug types and dosages. (**A**) Changes of the mean nuclear FOXM1-mVenus intensity over 48 h for different cell cycle perturbagens. n > 6000 cells per condition. (**B-C**) FOXM1 signals predict cell fate decisions. (B) Upper: Normalized FOXM1 activity for Palbociclib, Abemaciclib, BI2536, and Volasertib at 10 μM; A443654 at 1.25 μM, indicating peak delays (n > 6000 cells per condition). Lower: These drugs induce G1 arrest (n=3 per condition). (C) Upper: Normalized FOXM1 activity for CDK1/2III, RO3306, Volasertib, BI2536, and Danusertib at 1.25 μM; BMS265246 and Tozasertib at 5 μM, showing sustained FOXM1 increases at later time points (n > 6000 cells per condition). Lower: These drugs induce G2/M arrest (n =3 per condition).

### Single-cell analysis of FOXM1 dynamics reveals perturbagen-induced cell cycle delay or growth-arrested subpopulations

In our previous investigation, we identified distinct G1 and G2 arrest patterns in FOXM1 dynamics induced by 16 cell-cycle inhibitors across varying concentrations. However, a surprising finding emerged with ATR and CHK1 inhibitors, which exhibited FOXM1 dynamics and cell cycle distributions similar to the control group. This contrasted sharply with established expectations, as ATR inhibition typically induces truncated S phase and accelerated entry into the M phase (1). The limitations of bulk analysis became evident, failing to capture these nuanced effects (Fig. 5a). To address this, we employed single-cell tracking using the FUCCITrack method Taeib et al. (2022) (21) (Fig. S3a).

**Figure 5.**
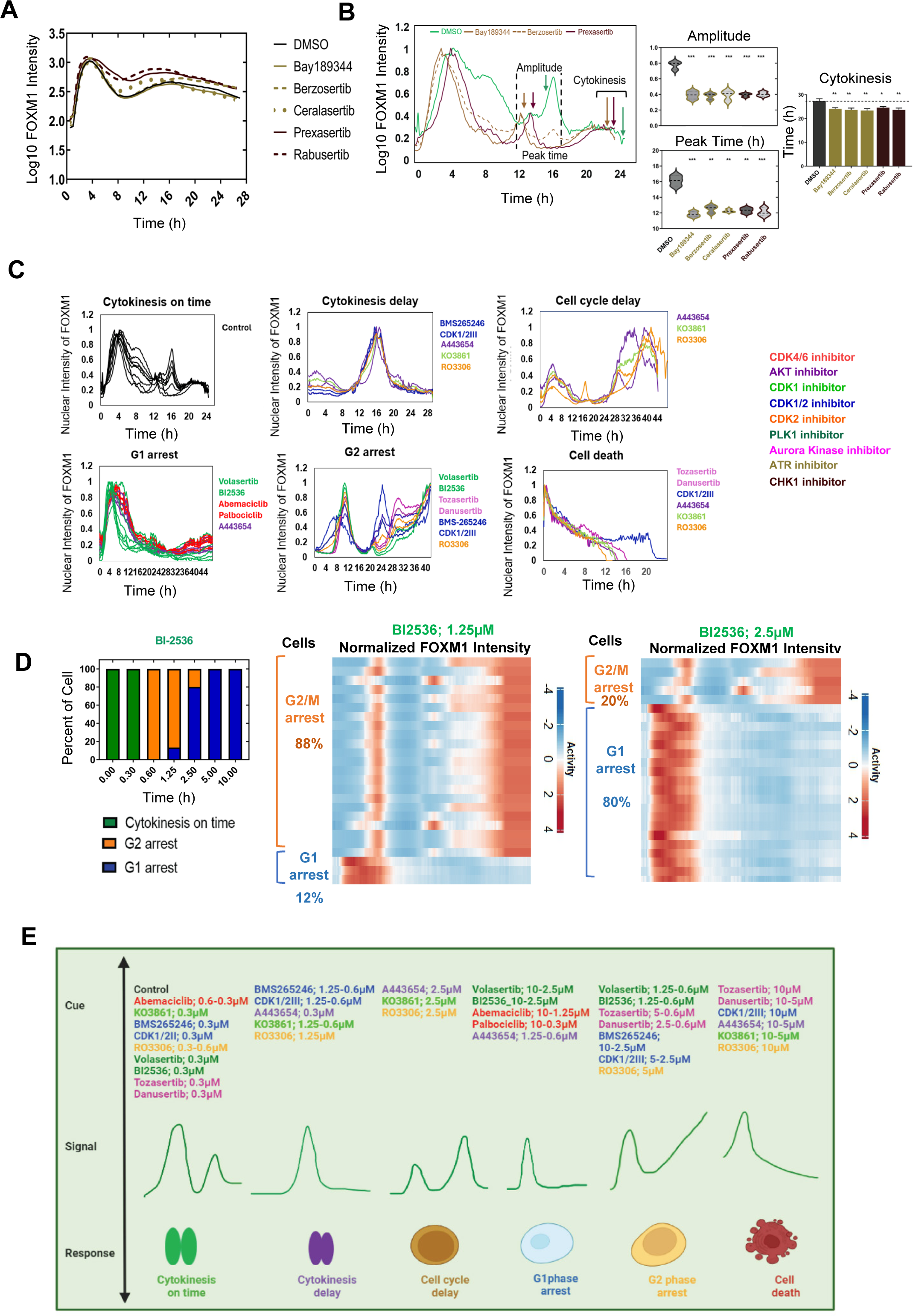
Single-cell FOXM1 dynamics are best predictive from individual cell fate decisions regardless of drug types and dosages. (**A**) Normalized FOXM1 activity for DMSO control (black), Bay189344 (olive line), Berzosertib (olive dashed), Ceralasertib (olive dotted), Presaxertib (maroon line), and Rabusertib (maroon dashed) at 10 μM plotted over 26 h (n > 6000 cells per condition). (**B**) Comparison of single-cell FOXM1 activity between DMSO, ATR (Bay189344, Berzosertib, and Ceralasertib), and CHK1 (Prexasertib and Rabusertib) inhibitors. Left: Single-cell trajectories post-treatment with DMSO (green), Bay189344 (olive), Berzosertib (olive dashed), and Prexasertib (maroon). The 2nd peak amplitude, peak time, and cytokinesis event are indicated with arrows. Middle: Distribution of FOXM1 2nd peak amplitude and peak time, compared between DMSO (black), ATR inhibitors (olive), and CHK1 inhibitors (maroon), based on single-cell traces from Fig. 5b-left (n=20 cells per condition). Right: Corresponding bar plot of cytokinesis events. *** p < 0.0005; ** p < 0.005; * p < 0.05. (**C**) Representative FOXM1 trajectories at single cells from 6 different phenotypic outcomes: (1) cytokinesis on time, (2) cytokinesis delay, (3) cell cycle delay, (4) G1 arrest, (5) G2 arrest, and (6) cell death. The colors of FOXM1 trajectories correspond with the inhibitor types. (**D**) Heterogeneity of phenotypic outcomes observed under BI2536 treatment. Left: Cell cycle distribution changes at varying drug concentrations (n = 25 cells per condition). Middle and Right: Heatmaps depicting single-cell FOXM1 activity at 1.25 μM (middle panel) and 2.5 μM (right panel) (n = 25 cells per condition). Ultimate phenotypic outcomes annotated on the left of each heatmap. (**E**) Summarizing schematic highlighting the cue-signal-response relationship from the different FOXM1 dynamical patterns from different drug types and concentrations.

Contrary to bulk analysis, single-cell tracking revealed distinct FOXM1 dynamics between DMSO and ATR/CHK1-treated cells, particularly in cytokinesis acceleration. FOXM1 pulse amplitude was 2-fold lower in ATR/CHK1-treated cells compared to DMSO (Fig. 5b-left panel), with peak times occurring approximately 4 hours earlier (median comparison) (Fig. 5b-middle panel). Moreover, ATR and CHK1 inhibitors accelerated cytokinesis by about 1.8 hours compared to DMSO (Fig. 5b-right panel), aligning with previous research and highlighting the complexities of cellular responses to these inhibitors. This refined analytical approach enabled us to uncover subtle yet significant shifts in cell cycle progression that were obscured by bulk analyses.

This observation makes us question whether traditional cell cycle analyses may have overlooked additional cell cycle phenotypes. We identified six distinct FOXM1 expression profiles (Fig. 5c and Fig. S3b): cells with a delayed first peak without a subsequent peak were in G1 phase arrest, while those with a continuous rise to a second peak indicated G2 arrest. A delayed first peak followed by a second peak indicated cell cycle delay, and cytokinesis delay was characterized by a much smaller first peak compared to the second. Cells undergoing cell death showed decreasing FOXM1 intensity over time (Fig. 5c-right panel).

### FOXM1 signals are predictive of cellular phenotype regardless of drug types and dosages

We postulated from the prior findings that FOXM1 signals could predict and explain the heterogeneous cell fate decision down to the single-cell resolution regardless of the cue type. Our analysis revealed a particularly striking example from PLK1 inhibitor (BI2536), where we observed a diverse range of FOXM1 dynamics leading to different cellular responses (Fig. 5d-middle and right panel). Specifically, we could categorize these phenotypic outcomes into three main groups, namely 1) cells undergoing G1 arrest, 2) cells entering G2 arrest, and 3) those completing cytokinesis in normal cell cycle time (Fig. 5d-left panel).

This pattern of heterogeneity was not exclusive to PLK1 inhibitors but also with Aurora kinase, CDK1/2, CDK4/6, and CDK1 treatments (Fig. S4a -d). Aurora kinase inhibitors induced cell death and G2 phase arrest in subsets of cells at concentrations as low as 5, 2.5, and 1.25 µM, correlating with FOXM1 activity (Fig. 5c and Fig. S4a). CDK1/2III inhibitor showed diverse responses, with cell death occurring at higher concentrations. Conversely, the CDK4/6 inhibitor Abemaciclib revealed a small subset of cells maintaining normal cell cycle dynamics amidst G1 arrest at 2.5 and 1.25 µM, while Palbociclib showed uniform effects (Fig. S4b, c). Interestingly, AKT and CDK2 inhibitors did not exhibit heterogeneity in their FOXM1 dynamics (Fig. S4e). These findings underscore extensive variability in cellular responses to identical stimuli at the single-cell level (Fig. 5e).

## DISCUSSION

We developed and characterized a FOXM1 reporter to track FOXM1 activity during cell cycle changes in MCF10A cells. Overexpression of FOXM1 typically induces Cyclin B1 and PLK1 expression(29, 30). To prevent such overexpression, we employed rigorous sorting steps, confirming that our reporter did not disrupt the dynamics of key FOXM1 target proteins like Cyclin B1 and PLK1 (Fig. 1h). This approach ensured our reporter system reliably captured intrinsic FOXM1 signaling dynamics without introducing confounding effects from downstream signaling alterations.

FOXM1 is known to be cell cycle-dependent (5, 30, 31), with its overexpression linked to various cancers (31-35). Different stages of cell cycle inhibition have distinct effects on FOXM1 expression: G1/S phase inhibition typically downregulates FOXM1, whereas G2/M phase inhibition often sustains or increases FOXM1 levels (6, 36). However, measuring how the dynamics of FOXM1 change in a perturbed cell cycle in real time is challenging. Our FOXM1 reporter addresses the challenge of real-time monitoring of FOXM1 dynamics during perturbed cell cycles by (1) enabling live microscopy-based quantification of FOXM1 activity, (2) providing high specificity to FOXM1, and (3) assessing how FOXM1 activity responds to diverse perturbations. Using this tool, we identified novel insights into FOXM1 dynamics under various cell cycle perturbagens. For instance, PLK1 inhibitors primarily affect the G2/M cell cycle phase (37), but can also induce G1 arrest through DNA damage response and p53 stabilization (37, 38). This result is consistent with our observation from the fPCA which showed that treating cells with a PLK1 inhibitor at 10 µM induced FOXM1 dynamics similar to Palbociclib, a CDK4/6 inhibitor, as confirmed by cell cycle analysis. In contrast, Abemaciclib, a Pan-CDK inhibitor targeting CDK2, CDK1, and others, affects both G1 and G2 phases distinctly (38). Our FOXM1 reporter effectively distinguished FOXM1 dynamics influenced by Abemaciclib and Palbociclib, revealing unique clustering patterns in fPCA analysis. This reporter enables real-time monitoring of FOXM1 dynamic patterns under diverse cell cycle perturbations, surpassing conventional exogenous expression techniques (6, 36).

The investigation of cell fate decisions has been further elucidated using live reporters. Specifically, studies have shown that live reporters enable researchers to monitor the dynamic changes of key intracellular signals linked to specific cell fate decisions at the single-cell resolution (17, 39-45). For instance, pulsatile P53 signaling can induce senescence, while cells experiencing large P53 pulses often undergo cell death (46-48). Similarly, our study revealed heterogeneous signal-to-phenotype relationships at single-cell resolution. High doses of PLK1 inhibitors (2.5-10 µM) shifted FOXM1 activity towards patterns resembling G1 arrest, whereas medium doses (0.6-1.25 µM) induced sustained increases leading to G2/M arrest. This highlights the critical role of dosing in shaping FOXM1 signals, with individual cell outcomes best predicted by FOXM1 activity dynamics. Our findings underscore the importance of analyzing dynamic signaling patterns over static biomarkers, aligning with the paradigm that cellular responses depend on temporal signal dynamics rather than the mere presence of cues(46, 47). Therefore, an invention of novel approaches to therapeutically targeting FOXM1 will require careful monitoring and perhaps the modulation of its signaling dynamics using combinatorial treatments of cell cycle inhibitors.

Cells can exhibit varied signaling dynamics (e.g., pulsatile, sustained high, sustained low) in response to the same perturbagen, influencing diverse cell fate decisions (49, 50). For example, Phongkitkarun et al., 2023, observed distinct cellular fates single-round or two-round division following TMP administration, linked to FOXM1 signaling dynamics (51). In our study, treatment with Aurora Kinase and PLK1 inhibitors at different doses also revealed heterogeneity in cellular responses. Aurora Kinase inhibitors induced G2/M arrest and occasional cell death, while PLK1 inhibitors caused cell populations to arrest either in G1 with delayed FOXM1 peak or in G2/M with sustained FOXM1 levels, likely due to delayed FOXM1 degradation (51). The observed heterogeneity in FOXM1 response highlights the importance of considering individual cell behaviors to understand drug efficacy and cellular outcomes.

Our study employs a biosensor-based approach to explore FOXM1 dynamics and investigate heterogeneous cellular responses to various cell cycle perturbations. The observed variability in cell responses emphasizes the necessity for personalized therapeutic strategies that precisely target FOXM1 activity. Factors such as stochastic gene expression (52), cell cycle asynchrony (47), and microenvironmental cues (53) likely contribute to these diverse FOXM1 responses. Understanding these factors is crucial for optimizing FOXM1-targeted therapies. While our study focused on established cell lines, future research should explore FOXM1 dynamics in patient-derived cancer models to elucidate clinical implications. Moreover, our study does not determine whether FOXM1 dynamics directly drive cellular responses or involve other mediators. Investigating how changes in FOXM1 dynamics affect downstream partners could provide further insights. Despite these limitations, our findings reveal a novel mechanism underlying cellular heterogeneity in drug responses, where intrinsic differences in FOXM1 signaling patterns among cells lead to varied phenotypic outcomes in response to cell cycle-modulating treatments.

## AUTHOR CONTRIBUTIONS

Conceptualization of the study: T.J, M.K, K.P, P.C and S.S. Methodology: T.J, M.K, K.P, P.C, S.S. Investigation and visualization: T.J. Formal analysis: T.J and M.K. Writing original draft, review and editing: T.J and S.S. Supervision: S.S. Resources: S.S. Funding acquisition: S.S.

## DATA AVAILABILITY STATEMENT

All data supporting the findings of this study are available within the paper and its supplementary information file.

## ETHICS APPROVAL STATEMENT

None

## IRB STATEMENT

None

