## Supplemental Figures for "Exploring the Single-Cell Dynamics of FOXM1 Under Cell Cycle Perturbations"

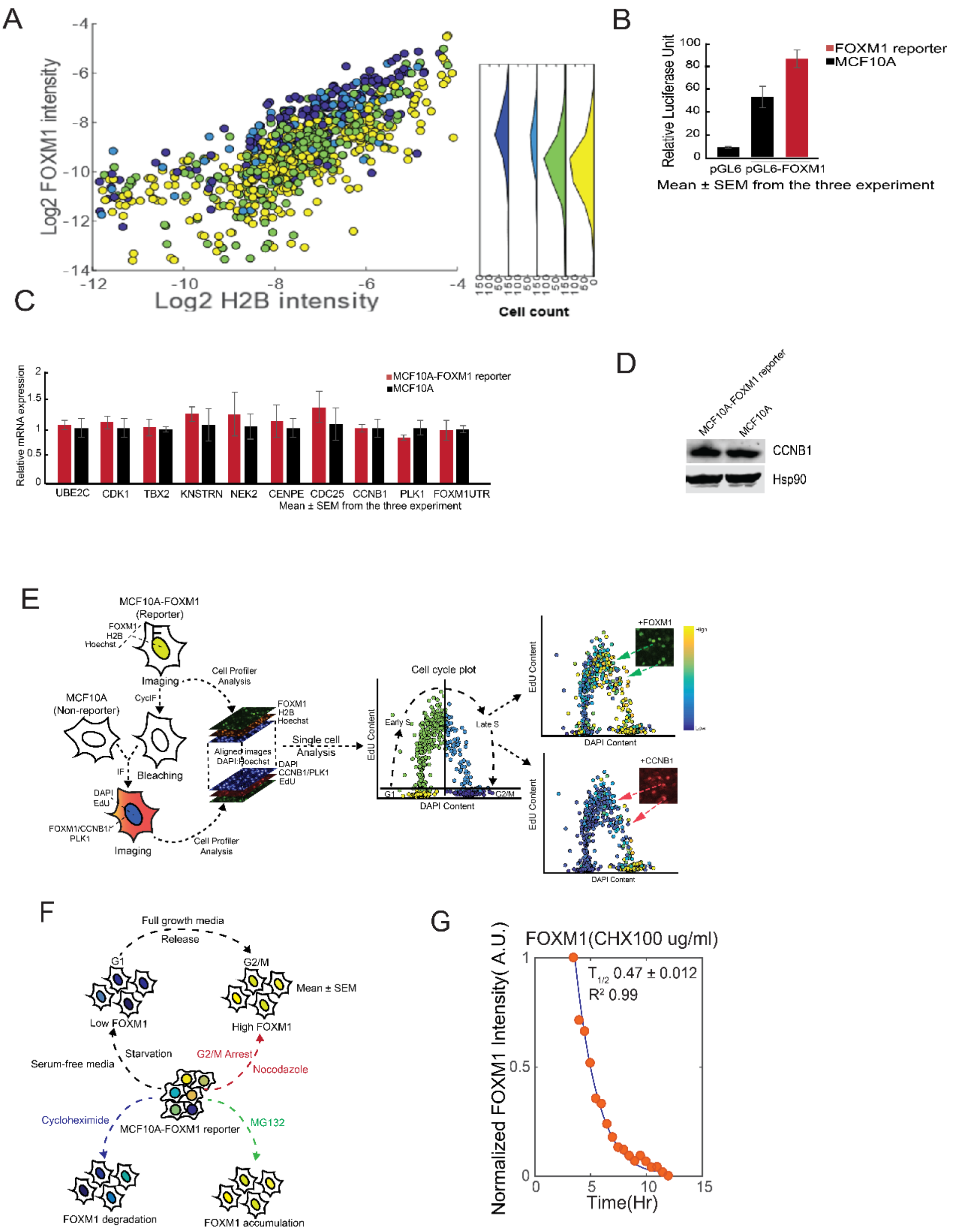


**Supplementary Figure 1. Comprehensive characterization of FOXM1-mVenus reporter activity and regulation in MCF10A Cells.** (A) Scattergram showing the landscape of reporter FOXM1 and mCherry single-cell activity. (B) Histogram showing fold induction of luciferase activity in MCF10A pGL6 and pGl6-FOXM1 and FOXM1 reporter. Data represent triplicate experiments ± SD. (C) mRNA expression of UBE2C, CDK1, TBX2, KNSTRN, NEK2, CNPE,

CDC25B, CCNB1, PLK1, and FOXM1 UTR in MCF10A cells that stably express FOXM1 sensor. Data were collected from synchronized cells in full growth media. (D) Western blot of CCNB1 in MCF10A wild type and MCF10-FOXM1 reporter. (E) Workflow showing the immunofluorescence staining. After serum-starve synchronization, cells were fixed and restained with EdU, DAPI, CCNB1, PLK1, and FOXM1 following the CycIF protocol. EdU incorporation and DNA content were used to mark cell cycle stages: G1, early S, late S, and G2/M. (F) Quantitative image-based analysis in cell cycle distribution and treatment after nocodazole treatment, cycloheximide or MG132. (G) Analysis of FOXM1 reporter degradation rate after cycloheximide treatment using high-content imaging.

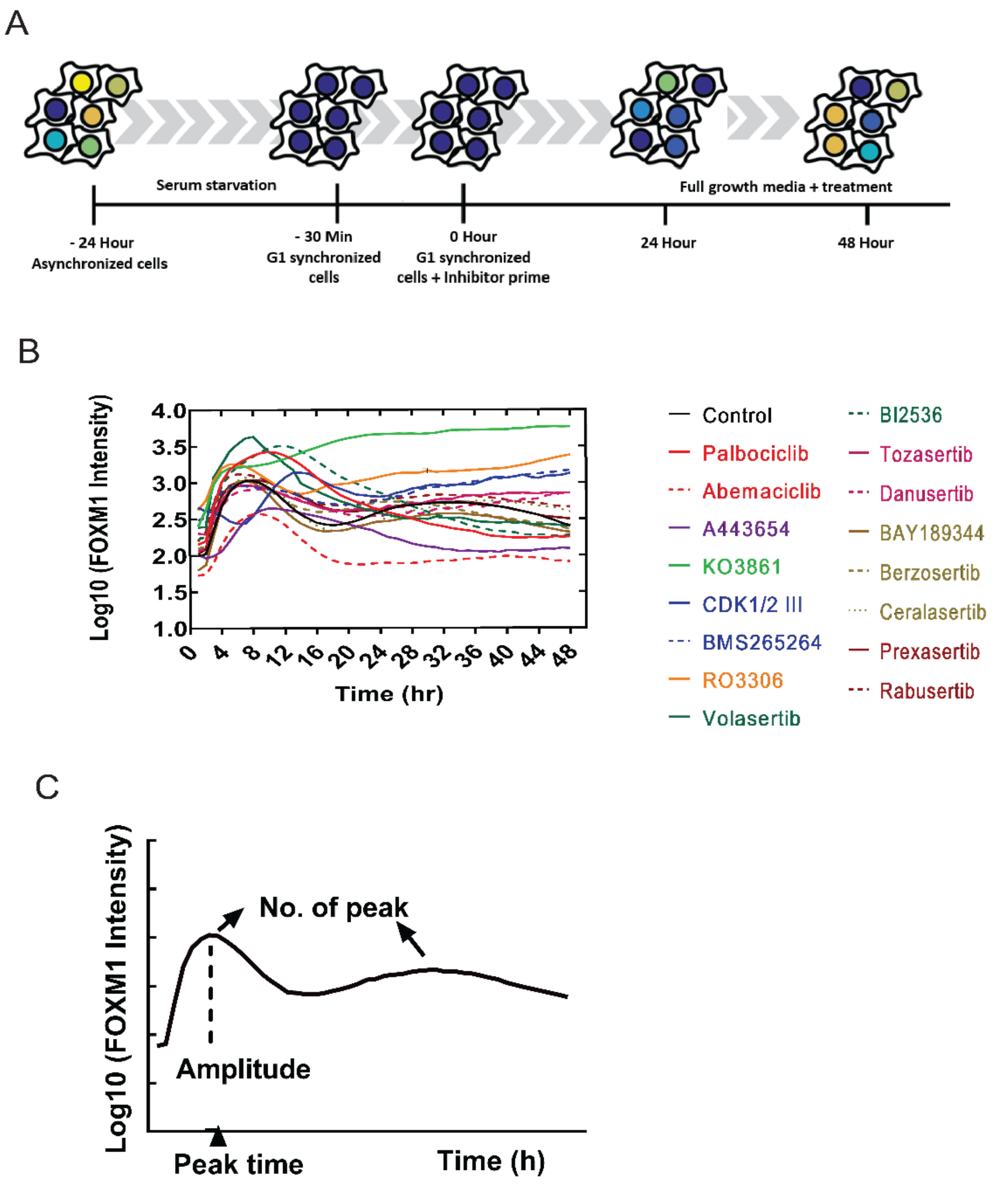


**Supplementary Figure 2. Analysis of FOXM1-mVenus reporter activity under cell cycle perturbations**. (A) Workflow for time-lapse imaging of FOXM1 reporter cells. FOXM1-mVenus reporter cells underwent 24 hours of serum starvation. 30 before serum replenishment, the cells were treated with a cell cycle inhibitor at concentrations ranging from 0 µM to 10 µM. Upon serum replenishment, the inhibitors were reintroduced at the same concentrations. Time-lapse imaging was then conducted for the subsequent 48 hours to observe the effects on the FOXM1-mVenus reporter cells. (B) Changes of the mean nuclear FOXM1-mVenus intensity over 48 h for different cell cycle perturbagens. n > 6000 cells per condition. (C) The scheme outlines the characterization of peaks.

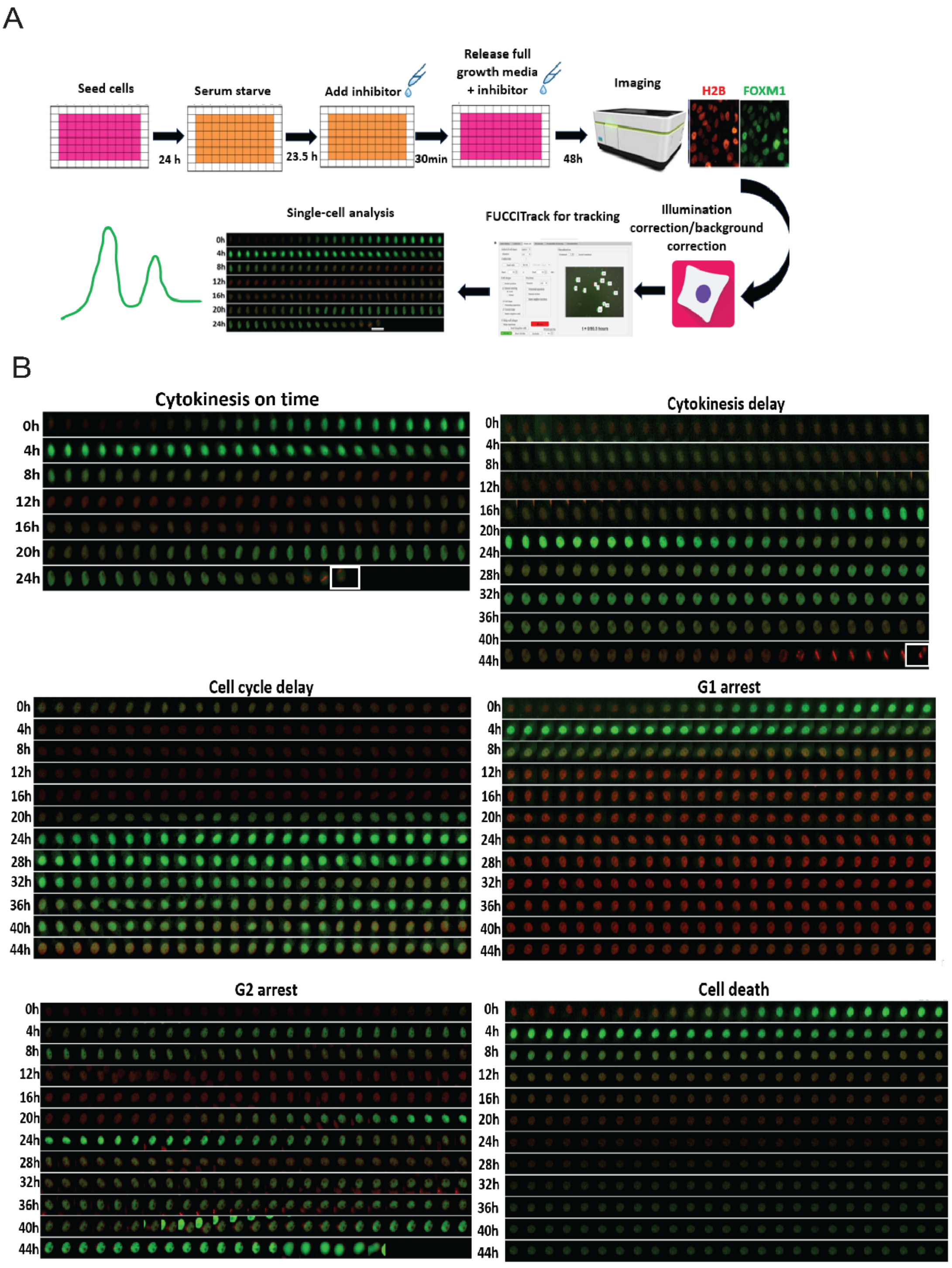


**Supplementary Figure 3. Single-cell tracking and phenotypic analysis of MCF10A-FOXM1-mVenus reporter cells.** (A) Scheme of single cell tracking MCF10A-FOXM1-mVenus reporter cells were serum starved for 24 h, then the inhibitor was added 30 min prior to the addition of full growth media. Later the cells were released in full growth media containing inhibitors. The cells were imaged every 10 min for 50 h. Background subtraction and illumination correction was performed through the cell profiler. FUCCITrack were used for cell tracking.

(B) Representative FOXM1 fluorescent microscopic images captured in different channels; FOXM1-mVenus (green) and H2B-mCherry (red) illustrating 6 different phenotypic outcomes: (1) cytokinesis on time, (2) cytokinesis delay, (3) cell cycle delay, (4) G1 arrest, (5) G2 arrest, and (6) cell death.

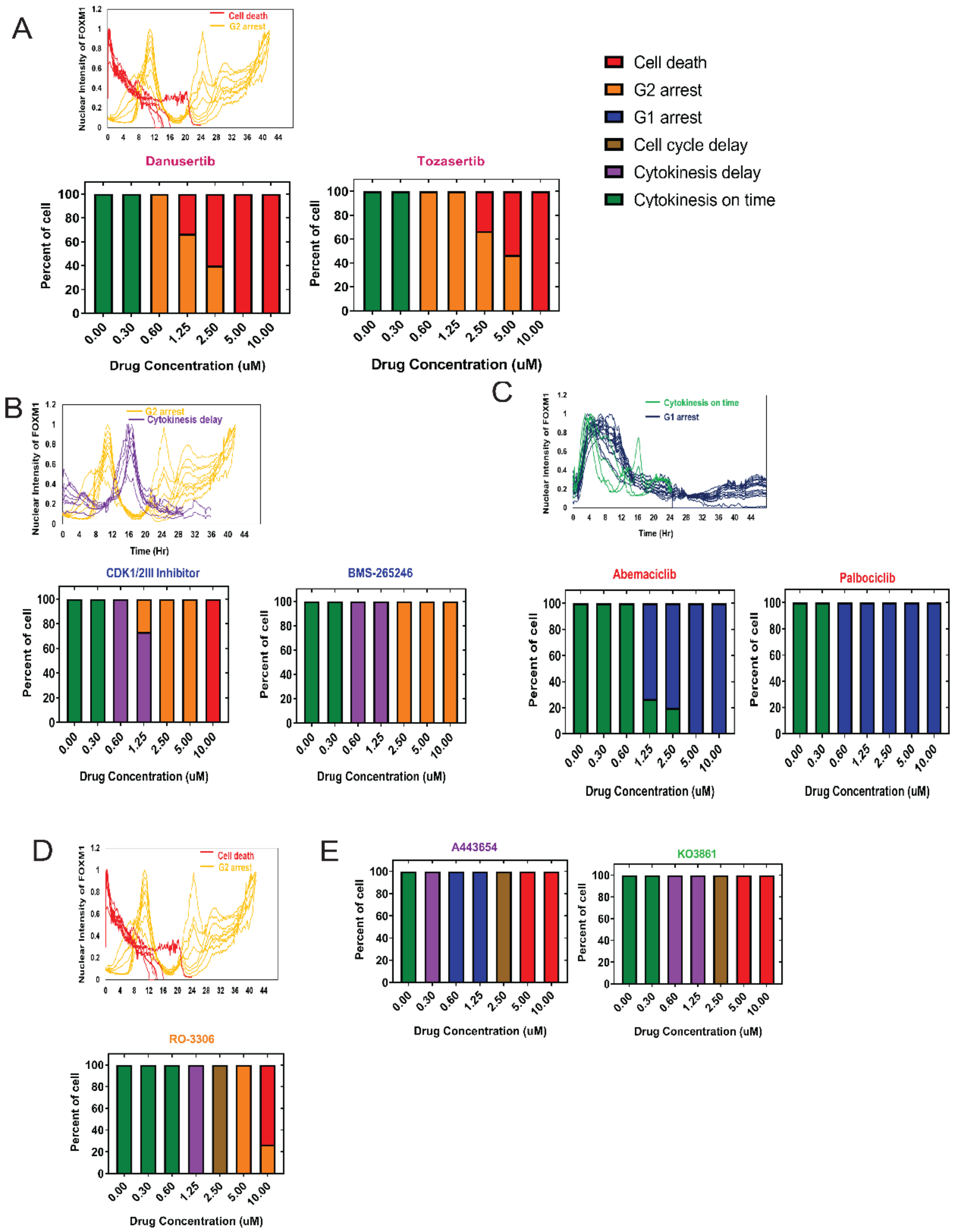


**Supplementary Figure 4. The phenotypic outcomes of FOXM1 reporter cells exhibit heterogeneity under varying cell cycle perturbations.** (A-D) Heterogeneity of phenotypic outcome at the individual cells could be observed from the same treatment condition. Cells were treated with cell cycle inhibitors from 0μM to 10 μM. Upper: Single-cell trajectories of FOXM1 reporter (n = 25 cells per condition). Lower: Cell cycle distribution changes at different drug concentrations (n = 25 cells per condition). (E) Cell cycle distribution changes at different drug concentrations (n = 25 cells per condition). Left: AKT (A443654) inhibitor. Right: CDK2 (KO3861) inhibitor.
